# Isolation of potential plant growth-promoting bacteria from nodules of legumes grown in arid Botswana soil

**DOI:** 10.1101/2020.09.02.257907

**Authors:** Melissa Kosty, Flora Pule-Meulenberg, Ethan A. Humm, Pilar Martínez-Hidalgo, Maskit Maymon, Sophia Mohammadi, Josh Cary, Paul Yang, Krisanavane Reddi, Marcel Huntemann, Alicia Clum, Brian Foster, Bryce Foster, Simon Roux, Krishnaveni Palaniappan, Neha Varghese, Supratim Mukherjee, T.B.K. Reddy, Chris Daum, Alex Copeland, Natalia N. Ivanova, Nikos C. Kyrpides, Tajana Glavina del Rio, Emiley A. Eloe-Fadrosh, Ann M. Hirsch

## Abstract

As the world population increases, improvements in crop growth and yield will be needed to meet rising food demands, especially in countries that have not developed agricultural practices optimized for their own soils and crops. In many African countries, farmers improve agricultural productivity by applying synthetic fertilizers and pesticides to crops, but their continued use over the years has had serious environmental consequences including air and water pollution as well as loss of soil fertility. To reduce the overuse of synthetic amendments, we are developing inocula for crops that are based on indigenous soil microbes, especially those that enhance plant growth and improve agricultural productivity in a sustainable manner. We first isolated environmental DNA from soil samples collected from an agricultural region to study the composition of the soil microbiomes and then used *Vigna unguiculata* (cowpea), an important legume crop in Botswana and other legumes as “trap” plants using the collected soil to induce nitrogen-fixing nodule formation. We have identified drought-tolerant bacteria from Botswana soils that stimulate plant growth; many are species of *Bacillus* and *Paenibacillus*. In contrast, the cowpea nodule microbiomes from plants grown in these soils house mainly rhizobia particularly *Bradyrhizobium*, but also *Methylobacterium* spp. Hence, the nodule microbiome is much more limited in non-rhizobial diversity compared to the soil microbiome, but also contains a number of potential pathogenic bacteria.

## Introduction

Botswana is a land-locked, semi-arid African country with a population of over 2 million people. The Kgalagadi (formerly Kalahari) Desert covers approximately 70% of Botswana. Temperatures reach as high as 38–46°C (110–115°F) and, coupled with an exceptionally low rainfall, this results in Botswana being described as “one of the most desertified countries in sub-Saharan Africa” [1]. Rainfall in the Kgalagadi Desert ranges from about <150 mm in southwestern Botswana to ca. 650 mm or more in the north (Okavango Delta) depending on the year. Botswana thus relies almost exclusively on imports for food and fuel, which ultimately is not sustainable.

Most of the agriculture in Botswana takes place in savannah ecosystems, defined as the lands between forested regions and deserts. Savannahs make up 10-25% of the world’s land surface [2], and in many parts of the world, including Africa, they have a dual purpose: growing crops and grazing livestock. The natural vegetation of savannah ecosystems in Botswana consists of grasses, thorny shrubs and trees, particularly vachellias (formerly known as acacias), as well as numerous native herbaceous dicotyledons, especially nitrogen-fixing legumes such as *Tephrosia purpurea* and *Indigofera charlieriana*. Many agricultural crops are cultivated in savannahs in Botswana, the most common being cereals, especially sorghum, sweet sorghum, millet, and maize. Other non-cereal crops include sunflowers, watermelons and nitrogen-fixing crops including beans, cowpea, and groundnut. Farmers are only able to meet 18% of the domestic demand for cereals [3], and crop production overall makes up a very small percentage (ca. 3.0%) of the agricultural gross domestic product (GDP) of Botswana. As a result, most food other than beef is imported from elsewhere.

Agricultural crop production in Botswana is challenging in large part due to the lack of reliable rainfall, coupled with a high evapotranspiration rate. Droughts have been known to last 15-20 years, with alternating periods of rainfall punctuated by drought years [4]. Furthermore, only 0.65-0.7% of Botswana is arable, and the sandy soils (arenosols) are highly susceptible to wind erosion. Arenosols are also frequently nutrient-poor, especially with regard to nitrogen, phosphorus, and potassium. Moreover, much of the soil is contaminated by pesticides and other toxic substances [5]. Like most of sub-Saharan Africa, the agricultural sector in Botswana is therefore extremely vulnerable to changes occurring worldwide in terms of climatic fluctuations and population growth.

Plant growth-promoting bacteria (PGPB) improve plant growth via diverse mechanisms including nitrogen fixation, phosphate solubilization, siderophore production for iron acquisition, as well as antagonizing pathogens, among others. The objective of this research was to assess microbial diversity in an agricultural soil in Botswana to identify rhizobial strains and PGPB, which are adapted to this ecosystem. To do this, we performed both 1) cultivation-independent analysis of the soil and also of legume root nodules to determine the differences in microbial diversity in addition to 2) cultivation-dependent analyses (i) of rhizospheres collected in two different years and (ii) isolations of plant-selected bacteria from the interior of root nodules of different legumes used as trap plants [6,7]. Because only 1-3% of bacteria in soil can be cultured, we reasoned that the nodule-isolated bacteria that could grow on artificial media might serve as potential inoculants because they were selected by the legume host.

Bacterial isolates from nodules and soil were both evaluated for their potential for PGPB activity using various assays. Specifically, we tested for bacteria that (i) promoted plant growth in N-limiting substrates, (ii) solubilized phosphate, (iii) produced siderophores, (iv) were salt and pH-tolerant, and (v) were not known pathogens. These traits (and others) strongly suggest plant growth promotion ability [6,7].

Even though the number of soil isolates that can be cultured is low compared to the number detected by cultivation-independent methods, many PGPB with potential for use in sustainable agriculture have been isolated from arid soils worldwide [7-10]. In addition, because legume nodules house bacteria in addition to rhizobia, we isolated non-rhizobial microbes from nodules because they are hypothesized to help in the establishment and maintenance of an effective nitrogen-fixing symbiosis [7]. Ultimately, the use of PGPB isolates from drylands for agricultural production will help reduce the amount of N fertilizer used in soils impacted by climate change and decrease agricultural runoff and fertilizer costs for Sub-Saharan farmers.

## Materials and Methods

### Soil collection and storage

Soil collections were made in 2017 and 2019 from the farm of the Botswana University of Agricultural and Natural Resources in Notwane (coordinates 24° 34’ 48.5” S, 25° 57’ 46.7” E, elevation 973 m) from under the canopy of the indigenous legume plant *Tephrosia purpurea* (Figure S1). *Lablab purpureus* was growing in close proximity to the site. A preliminary collection made in 2013 predicted that the soil microbial diversity was quite high (data not shown). The general vegetation of this region is mixed savannah, and the soils are deep Kalahari sands (Arenosols according to the FAO classification) with no soil horizons and very little humus [11]. The mean annual rainfall is about 300-400 mm per annum. A trowel of soil was collected into a clean plastic bucket from the rhizosphere of four randomly selected *T. purpurea* plants in an area of about 5 × 5 m. The sample was collected from a depth of between 10 to 15 cm from the soil surface. The collected soil was brought to the laboratory at BUAN where it was thoroughly homogenized in a flat plastic tray and subdivided into four quarters. Two of the quarters were discarded and the procedure was repeated until the required amount of approximately 100 g was obtained. Soil was sent to the UCLA laboratory under USDA permit P526P-19-03102. The 2017 and 2019-collected soils were analyzed in January 2019 (Table 1). All soil samples were stored at 4°C after collection.

**Table 1.**
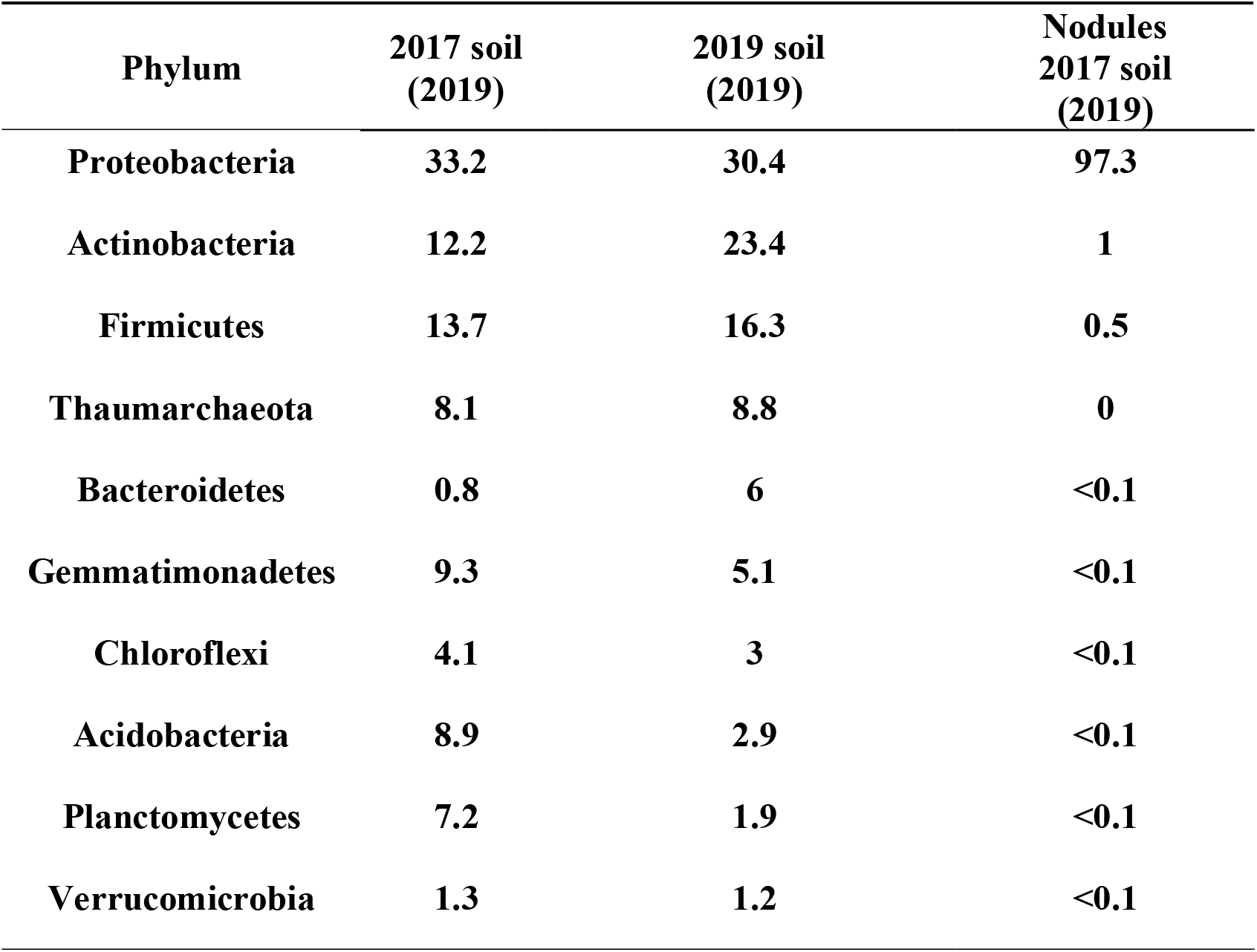
Percent relative abundance of bacterial phyla in soil samples and cowpea nodules. The year in parentheses indicates the date that the sample was analyzed.

### Soil environmental DNA (eDNA) extraction, PCR amplification, and pyrosequencing

eDNA was extracted from each collected soil sample, starting with a sample size of 0.25-0.5 g per isolation. Two PowerBead preparations of 0.5 g were used along with the PowerSoil Pro DNA Isolation kit (MoBio Labs/QIAGEN, Carlsbad, CA, USA) according to the instructions. The samples were combined during the first MB Spin Column step (step 8). The quality of extracted DNA was assessed by visualization on a 1% agarose gel with a high molecular mass ladder (Invitrogen, ThermoFisher Scientific, LLC, USA).

A commercial service (MR DNA, Shallowater, TX) performed *rrs* gene sequencing, which was analyzed using the Illumina platform. Pipeline data in OTU form were processed with Krona tools in Excel to generate data and visualized to describe the relative abundance of bacterial classes in the sample. Additional sequencing was performed by the DOE Joint Genome Institute (USA).

### Isolation of bacteria from soil

Serial dilutions of soil were made and samples from the 10^−2^ – 10^−6^ dilutions were plated onto a number of culture media, including Luria-Bertani (LB), Reasoner’s 2A (R2A), Tryptone Yeast (TY), Yeast Mannitol Broth (YMB), and Arabinose Gluconic Acid (AG) agar media.

### Plant tests

Several legumes were tested initially for their responses to Botswana soil (Table 2), including a number of African crop plants, namely cowpea (*Vigna unguiculata* subsp. *unguiculata*, USDA accession PI339603 from South Africa) and Bambara groundnut (*V. subterraneana* Landrace Tanzania) [12]. Other legumes tested include *Lebeckia ambigua* (a South African native perennial forage legume) [13]; siratro (*Macroptilium atropurpureum*), a broad host range legume [14]; hyacinth bean (*Lablab purpureus*), an indigenous legume growing in the collection site [15]; and finally *Tephrosia virginiana* (goat’s rue) [16], which was used as a substitute for *T. purpurea* because the latter’s seeds were not available to us.

**Table 2.** Overview of different legumes inoculated with Botswana soil in nitrogen-deprived conditions, their sources, and their responses compared to non-inoculated nitrogen-deprived control plants.

| Legume Species | Soil Tested | Nodulation Response<br>+ or - | Place of Origin | Source of Seed | Ref. |
| --- | --- | --- | --- | --- | --- |
| *Vigna unguiculata* subsp. *unguiculata* (Cowpea) | 2017 & 2019 | + | South Africa | US Department of Agriculture accession PI339603 | [12] |
| *V. subterranea* (Bambara groundnut) | 2019 | + | Tanzania | Felix Dakora, Tshwane University, Pretoria, South Africa | [12] |
| *Lebeckia ambigua* | 2019 | - | Fynbos, Western Cape, South Africa | Murdoch University, Perth, Australia | [13] |
| *Macroptilium atropurpureum* (Siratro) | 2019 | + | Tropical and subtropical America | Bio-Next, Wichita, KS | [14] |
| *Lablab purpureus* (Hyacinth bean) | 2019 | - | Africa | Onalee Seeds, LLC, Madeira Beach, FL 33738 | [15] |
| *Tephrosia virginiana* (Goat's rue) | 2019 | + | Eastern US and Canada | Prairie Moon Nursery, Winona MN 55987 | [16] |

Legume seeds were surface-sterilized in ethanol for 1-2 min. and then in different percentages of bleach depending on the seed type. *L. ambigua, T. virginiana*, and siratro seeds were scarified for 1 min. prior to surface sterilization. After the sterilization procedure, the seeds were washed extensively in sterile water and planted in 2% water agar plates until the seeds germinated. The seeds were then transferred to 3.8-liter autoclavable pots filled with a 2:1 mixture of vermiculite and perlite. Prior to planting, both the pots and the vermiculite and perlite were sterilized by autoclaving (for 3 and 20-minute dry cycles, respectively).

Each trap experiment consisted of three treatments: a nitrogen-deprived experimental group inoculated with Botswana soil, and two uninoculated control groups either treated with nitrogen (Control +N) or deprived of nitrogen (Control -N). For experimental plants, 10 g of the soil to be tested were mixed in with the vermiculite and perlite. Soils stored for a long time were incubated in 5 ml of water overnight at 30°C before addition to the pots. Plants were grown in a Conviron growth chamber or in a greenhouse within the UCLA Plant Growth Center. Control plants provided with nitrogen were watered weekly with quarter-strength Hoagland’s medium (per liter: 2 mL 1 M MgSO_4_; 1 mL 1 M KH_2_PO_4_; 1 mL FeEDTA stock; 1 mL micronutrients stock; 625 μL 1M Ca(NO_3_)_2_; 625 μL 1M KNO_3_; pH 5.7-5.8.) The experimental and nitrogen-deprived control groups were watered with modified Hoagland’s medium as follows: for the first 6-8 weeks they were watered with 25% N (the amount of Ca(NO_3_)_2_ and KNO_3_ was reduced to 156.25 μL per liter), and then 0 % N until they were harvested at 9-12 weeks.

Two separate major cowpea trap experiments were performed: the first with the 2017 soil sample, and a second using the 2019 sample. For each experiment, a total of 12 control -N, 12 control +N, and 24 experimental cowpea plants were planted initially (three seeds per pot). Plants were photographed and dry weights were calculated; statistical significance of measurements was assessed via one-way ANOVA and Tukey’s post-hoc test.

### Isolation of bacteria from root nodules

Green, healthy plants were removed from the experimental treatments and the nodules were collected at different time points, but usually after 10 weeks. Bacteria were isolated from surface-sterilized root nodules of the trap legume plants that nodulated. The nodules were surface-sterilized (70% ethanol for 1 min. followed by 10% commercial bleach for 10 min.) and rinsed numerous times with sterile water. Nodules were then squashed with a sterile glass rod, and the crushed nodule suspension was diluted in sterile distilled water and dilutions were plated out on various agar media, including Luria-Bertani (LB), Reasoner’s 2A (R2A), Tryptone Yeast (TY), Yeast Mannitol Broth (YMB), and Arabinose Gluconic Acid (AG) agar media. Distinct colony morphologies were chosen from the different plates, and the bacteria were plate-purified and stored as glycerol stocks at −80°C. Liquid bacterial cultures were grown in LB, TY, AG, or YMB medium overnight. For solidified media, 15 g of bacterial agar (Sigma) were added per 1 liter of culture and the bacteria were cultured overnight or longer.

### Tests for PGPB activity, salinity, and pH tolerance

After several rounds of plate purification, soil and nodule-isolated bacteria were tested for their ability to solubilize CaH_2_-phosphate (modified Pikovskaya medium; PVK [17]), produce siderophores (Chrome Azurol S; CAS [18]), and hydrolyze cellulose (CMC [19]). *Paraburkholderia unamae* MTI-671^T^, a nitrogen-fixing, beta-rhizobial strain isolated from Mexico [20] was used as a positive control for the PVK assay and *Bacillus subtilis* 30VD-1 [21] was used as a control for the CAS and CMC plates. Isolates were also tested for the ability to produce amylase by cultivation on starch plates for 48 h, after which the plates were flooded with Gram’s iodine.

In addition, the isolates were grown on LB or TY plates containing no additional NaCl as well as on plates containing 3%, 4%, and 5% salt for one week. For testing pH tolerance, LB or TY plates at pH 5, 7, and 9 were used to grow the isolated bacteria (Table 4). Positive results on salinity and pH tolerance tests were evaluated by growth after one week in comparison to the no-salt and pH 7 controls.

**Table 3.** Cumulative list of bacteria isolated from nodules of cowpea plants grown as trap plants in *Tephrosia* soil.

| Closest Bacterial Identity | Family | Soil Sample | Known Interactions/Functions | Ref. |
| --- | --- | --- | --- | --- |
| *Rhizobium pusense* LMG 25623 | *Rhizobiaceae* | 2019 | Induces nitrogen-fixing nodules on *Cicer arietinum* L. | [40] |
| *Rhizobium tropici* SARCC-RB1h | *Rhizobiaceae* | 2017 | Induces nitrogen-fixing nodules on a diversity of legumes | [41] |
| *Rhizobium miluonense* Kav-b 4 / CCBAU4125<sup>T</sup> | *Rhizobiaceae* | 2017 | Induces nitrogen-fixing nodules on *Lespedeza* spp. and *P. vulgaris* | [42] |
| *Bradyrhizobium jicamae* PAC68<sup>T</sup> | *Rhizobiaceae* | 2019 | Induces nitrogen-fixing nodules on *Pachyrhizus* and *Lespedeza* spp. | [43] |
| *Bradyrhizobium arachidis* LMG 26795 | *Rhizobiaceae* | 2019 | Induces nitrogen-fixing nodules on *Arachis hypogaea* and *Lablab purpureus* | [44] |
| *Bradyrhizobium yuanmingense* CBMCC 1.3531 | *Rhizobiaceae* | 2019 | Induces nitrogen-fixing nodules on *Lespedeza* spp, cowpea, and *Glycyrrhiza uralensis*. | [45] |
| *Shinella yambaruensis* MS4 | *Rhizobiaceae* | 2019 | Utilizes 3-methyl sulfolane as sole sulfur source | [46] |
| *Methylobacterium dankookense* SW08-7 / E111CS6 | *Methylobacteriaceae* | 2017 | Originally isolated from drinking water | [47] |
| *Methylobacterium radiotolerans* JCM 2831 | *Methylobacteriaceae* | 2019 | Phytohormone and ACC deaminase production, possible nodulator | [48] |
| *Methylobacterium populi* BJ1001 | *Methylobacteriaceae* | 2019 | Aromatic compound degrader, phytohormone production | [49] |
| *Ochrobactrum anthropi* HK8 | *Brucellaceae* | 2017 | Emerging opportunistic pathogen | [50] |
| *Sphingomonas leidyi*<br>ATCC 15260 | *Sphingomonadaceae* | 2019 | Isolated from millipede hind gut | [51] |
| *Novosphingobium* sp.<br>THN1 | *Sphingomonadaceae* | 2019 | Degrades microcystin-LR | [52] |
| *Burkholderia dolosa*<br>FDAARGOS_562 | *Burkholderiaceae* | 2017 | Potential human pathogen | [53] |
| *Ralstonia mannitolytica* NK20 | *Burkholderiaceae* | 2017 | Emerging opportunistic pathogen | [54] |
| *Lysobacter oryzae*<br>YC6269<sup>T</sup> | *Xanthomonadaceae* | 2019 | Biocontrol activity against fungi | [55] |
| *Microbacterium lacusdiani* IXI CY01 | *Microbacteriaceae* | 2019 | Phosphate-solubilizing bacterium | [56] |
| *Staphylococcus pasteurii* A63 | *Staphylococcaceae* | 2017 | Potential human pathogen | [57] |
| *Bacillus nealsonii*<br>16BCA / Marseille-P2085 | *Bacillaceae* | 2017 | *Nicotiana attenuata* root endophyte | [58] |
| *Fictibacillus phosphorivorans*<br>1278 C12F/CA7<sup>T</sup> | *Bacillaceae* | 2017 | Phosphate-solubilizing bacterium | [59] |
| *Paenibacillus lautus*<br>LMG 11157 | *Paenibacillaceae* | 2017 | Opportunistic human pathogen | [60] |

**Table 4.** Characterization of bacteria isolated from root nodules of cowpea plants grown in the 2017 and 2019 Botswana soil samples. Tests include pH tolerance (pH 5 – 9), salinity tolerance (3-5% NaCl), siderophore production (CAS), amylase production (Starch), and phosphate solubilization (PVK).

| Closest<br>Bacterial Identity (%) | pH 5 | pH 7 | pH 9 | 3%<br>NaCl | 4%<br>NaCl | 5%<br>NaCl | CAS | Starch | PVK |
| --- | --- | --- | --- | --- | --- | --- | --- | --- | --- |
| *Bradyrhizobium jicamae*<br>PAC68 (100) | + | + | + | - | - | - | ng<sup>1</sup> | - | + |
| *Bradyrhizobium sp.*<br>AZCC_0077 (100) | + | + | + | - | - | - | ng | - | - |
| *Rhizobium milunoense*<br>Kav-b 4 / CCBAU4125T<br>(100) | + | + | + | - | - | - | + | - | + |
| *Rhizobium pusense* LMG<br>25623 (100) | + | + | + | + | + | - | + | - | + |
| *Rhizobium tropici*<br>SARCC-RB1h (100) | + | + | + | - | - | - | + | - | + |
| *Shinella yambaruensis*<br>MS4 (98.11) | + | + | + | - | - | - | ng | - | + |
| *Methylobacterium*<br>*dankookense* E111CS6<br>(98.68) | + | + | + | - | - | - | ng | - | - |
| *Methylobacterium*<br>*populi* BJ001 (99.76) | + | + | + | - | - | - | ng | - | - |
| *Methylobacterium*<br>*radiotolerans* JCM 2831<br>(99.76) | + | + | + | - | - | - | ng | - | - |
| *Sphingomonas leidy*<br>ATCC 15260 (99.45) | + | + | + | - | - | - | ng | - | - |
| *Novosphingobium sp.*<br>THN1 (99.54) | + | + | + | - | - | - | ng | - | - |
| *Lysobacter oryzae*<br>YC6269 (98.69) | - | + | + | - | - | - | ng | - | - |
| *Microbacterium*<br>*lacusdiani* JXJ CY 01<br>(99.87) | + | + | + | + | + | + | ng | + | - |
| *Bacillus nealsonii*<br>16BCA/Marseille-P2085<br>(100) | + | + | + | + | + | + | ng | - | - |
| *Fictibacillus*<br>*phosphorivorans* 1278<br>C12F/CA7<sup>T</sup> (100) | + | + | + | + | + | + | ng | + | - |

<sup>1</sup>ng, no growth.

### Genomic DNA extraction and bacterial isolate identification

Genomic DNA was extracted as follows: bacterial cells from a pure culture obtained from a single plate-purified colony were lysed by boiling at 100°C for 20 min in a 1:1 mixture of 0.1M NaOH and 0.5% SDS [9]. Samples were then cooled at −20°C for 10 min and centrifuged at 10,000 x g for 5 min.

The *rrs* genes were amplified as follows. One µl of DNA template was mixed with a PCR master mix (per 25 µl reaction, 17.5 µl H_2_O, 2.5 µl 10X PCR buffer with MgCl_2_, 2.5 µl 20 mM dNTP, 1 µl each of 2 mM forward primer fD1 and reverse primer rD1 [22], and 0.5 µl Taq polymerase.) The initial denaturation was for 5 min at 95 °C followed by a three-step cycle repeated 35 times (denaturation, 30 sec at 95 °C; annealing, 30 sec at 55 °C; extension, 60 sec at 72 °C,) and the final extension was for 10 min at 72 °C. The purified PCR products were sent for sequencing to Laragen (USA).

### Biosafety Tests

Because many plant and animal pathogens have been isolated from soil [23], the isolates were first identified by *rrs* gene analysis and any potential pathogens were discarded. Those isolates not immediately determined as pathogenic were tested in other assays [24] to verify their status as BSL1 (Biosafety Level 1) before they were used experimentally.

### Nodule environmental DNA (eDNA) extraction

eDNA was extracted from 0.5 g of squashed, surface-sterilized cowpea root nodules using the PowerSoil Pro DNA Isolation kit (MoBio Labs/QIAGEN, Carlsbad, CA, USA).

## Results and Discussion

Because only a small percentage of soil microorganisms can be cultured in the laboratory, we first undertook a cultivation-independent approach to obtain an overview of the diversity of microbes in Botswana soils in addition to using cultivation-dependent methods to discover nodule tissue-isolated microbes that might serve as inoculants for crops grown in dry, nutrient-poor soils. This two-pronged approach of using both cultivation-independent and -dependent methods provided an overview of the Botswana arid soil environment and yielded insight into microbial diversity. Due to the fact that only a very small number of microbes can be reliably cultured for further study towards using them as legume crop inoculants, we focused on the PGPB that the plant specifically selects within its root nodules with the goal of finding “helpers” for the nitrogen-fixing rhizobia in supporting plant growth.

### Cultivation-Independent Methods: Soil Microbiomes

Soil collections were made in 2017 and 2019 from the farm of the Botswana University of Agricultural and Natural Resources in Notwane from underneath an indigenous *Tephrosia purpurea* plant (Figure S1). Environmental DNA was isolated from both samples in 2019, at the same time trap experiments were performed, to provide insight into the diversity of the soil microbial community. It should be noted that there was a two-year gap between collecting the 2017 sample (during which time it was stored at 4°C) and its analysis in contrast to the 2019 sample that was analyzed almost immediately after collection. Because the methodology for DNA extraction was the same, we hypothesized that differences in the percentages of the phyla might occur from changes brought about by storage conditions, or time elapsed.

In both the 2017 and 2019 samples, the major phyla were Proteobacteria, Firmicutes, and Actinobacteria. The percentage of Actinobacteria in the 2019 soil was almost twice that of the 2017-collected soil (Figure 1, Table 1). These results are in line with those obtained by other authors. The bacterial genera responsible for the induction of N_2_-fixing nodules in legumes belong to the phylum Proteobacteria and are therefore part of the dominant group. The phyla Actinobacteria and Firmicutes contain several genera of bacteria with PGPB activities that are very well documented [7, 21].

**Figure 1.**
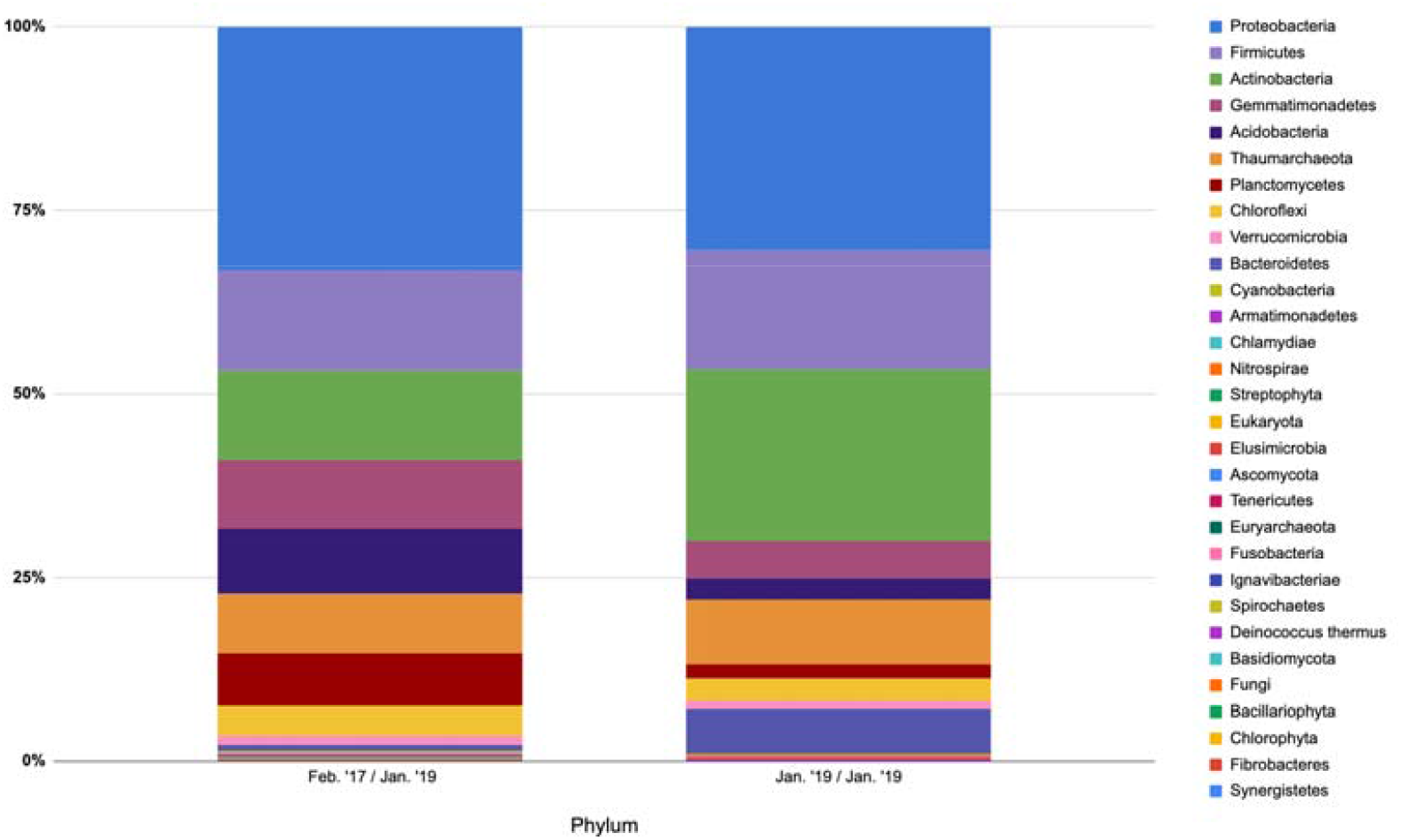
Botswana soil microbial community composition (by phylum) as determined by analysis of environmental DNA. The date on the left below each column indicates the date the soil sample was collected, whereas the right date indicates when the eDNA was isolated from the sample and analyzed.

The percentages of Gemmatimonadetes, Acidobacteria, and Planctomycetes also varied between the 2017- and 2019-analyzed samples (Table 1). Whether or not these differences are due to the delay between collection and analysis or other factors such as changes in the surrounding environment such as water content is not known. Nevertheless, the data demonstrated that the dominant microbes from the eDNA analysis were Proteobacteria, Firmicutes, and Actinobacteria, all of which are more likely to be cultured and serve as inoculants than the other bacteria listed.

### Cultivation-Dependent Methods

Soil isolates are often considered as sources of inoculants for crops in agriculture, particularly rhizobia and other plant growth-promoting bacteria (PGPB), but the nodule isolates may be a more specific inoculant for because they are found within nitrogen-fixing nodules [7]. Evidence based on coinoculation experiments with rhizobia [21, 25, 26] also indicates that soil-isolated as well as nodule-associated bacteria may be important for improving plant growth via plant nutrition. Although a large number of soil isolates have been tested for their ability to produce siderophores, solubilize phosphate, fix nitrogen, or perform other plant-growth promoting functions, to our knowledge only a few of them have been actively incorporated into agricultural practices. Due to the sheer numbers of soil isolates potentially available in Botswana soils (some of which are listed in Table S1), we focused our study on microbes housed in legume nodules.

Several trap plants, including *Vigna unguiculata* (cowpea), *Macroptilium atropurpureum* (siratro), and *Tephrosia virginiana*, nodulated following inoculation with Botswana soil mixed with an artificial substrate watered with -N medium, but cowpea gave the most consistent results (Table 2, Figure 2). Bacteria isolated from cowpea nodules included rhizobia (such as *R. tropici, Bradyrhizobium arachidis*, and *Microvirga* spp.), which are known to nodulate cowpea and other legumes [12]. Furthermore, species of *Bacillus*, including *B. safensis* and *B. pumilus*, well-known PGPB, were also isolated from cowpea nodules (Table 3). In addition, several possible opportunistic pathogens including *Ochrobactrum anthropi, Burkholderia dolosa, Ralstonia mannitolytica, Staphylococcus pasteuri* and others were isolated from cowpea nodules and identified by *rrs* sequencing. These emerging pathogens, which are often found in plant rhizospheres [23], were discarded. Non-pathogenic isolates were tested for PGP traits and their ability to grow under salinity stress and at different pH values (Table 4). A number of isolates exhibited possible PGP activity including phosphate solubilization and siderophore production.

**Figure 2.**
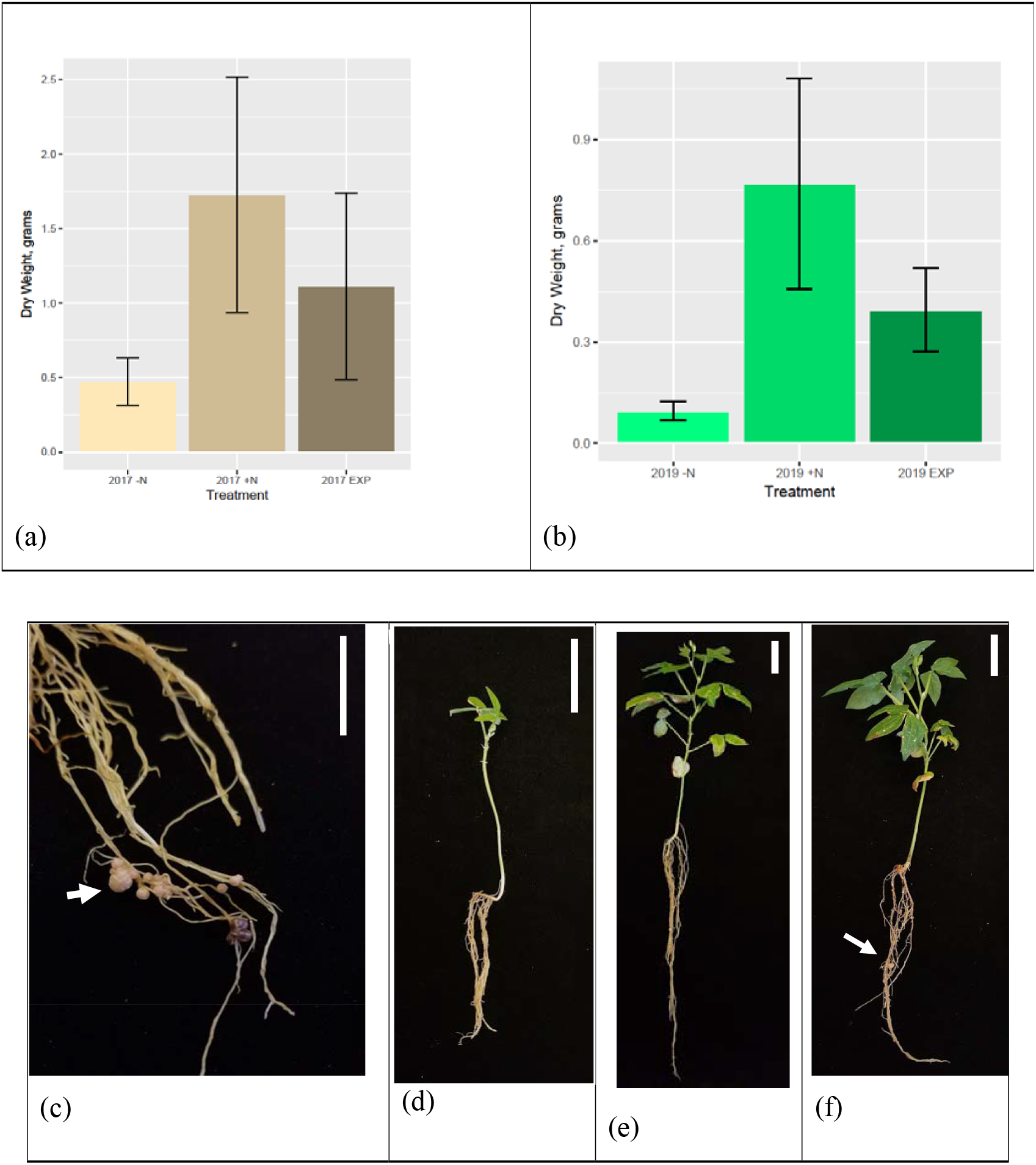
Trap experiments with cowpea plants grown in 2017 and 2019 Botswana soil samples. **(a)** Bar chart showing distribution of dry weights of nitrogen-deprived control plants (-N, n = 11), control plants grown with nitrogen (+N, n =10), and experimental plants grown in 2017 Botswana soil (EXP, n = 22). All groups were statistically different at a significance level of 0.05. Error bars show standard deviation. **(b)** Bar chart showing dry weights of cowpea plants grown in 2019 Botswana soil (-N, n=12; +N, n=10; EXP, n=17). All groups were statistically different at a significance level of 0.05. Error bars show standard deviation. **(c)** Nodules (arrows) on the roots of an experimental cowpea plant grown in 2019 soil, **(d, e, f)** Whole plant view of **(d)** control -N, **(e)** control +N, and **(f)** experimental cowpea plants grown in 2019 soil. Arrows point to nodules. Scale bar: 5 cm.

### Cowpea Trap Experiments

Cowpea plants grown in the 2017 Botswana soil sample were harvested after 9 weeks of growth. Control +N plants produced more biomass as measured by dry weight than plants from all other treatments, averaging 1.73 g. The experimental plants were darker green in color than control -N plants and produced more than twice as much biomass, averaging 1.11 g compared to 0.47 g for the -N control. Cowpea plants grown in the 2019 Botswana soil sample were harvested after 12 weeks of growth because of a lag in growth at the start. Control +N plants were larger, more robust, and darker green than experimental or control -N plants, averaging 0.77 g. Although the experimental plants were not as robust as the +N control plants, a result frequently observed in control plants given super-optimal N, the inoculated cowpeas produced significantly more biomass (0.396 g versus 0.096 g) than the -N controls. All experimental cowpea plants from both soil treatments developed multiple, pink-colored root nodules, whereas control -N and control +N plants were devoid of nodules. In both experiments, control -N plants were indistinguishable from control plants grown in soil that was sterilized by autoclaving (not shown) (Figure 2).

### Cultivation-Independent Analysis of Cowpea Nodule Microbiomes

Because cultivation methods are biased for the reason that very few bacteria are capable of growing on standard bacteriological culture media, we analyzed the cowpea nodule microbiome by isolating eDNA from the nodule tissue and sequenced the eDNA with the goal of obtaining an inventory of the nodule microbial population. We predicted that these analyses would give us insight into the bacteria that were specifically selected by the plant and if they were culturable, they might have potential to be used as commercial inocula.

As expected from anatomical studies of determinate nodules such as cowpea [27], the nodule interior based on eDNA analysis is dominated by *Bradyrhizobium* spp. (Figure 3). Although DNA sequences from numerous bacterial genera including *Microvirga, Rhizobium, Bacillus, Sphingomonas*, and others were detected in the nodule microbiome in this study (Table 3), the exact percentages and diversity of non-rhizobial microbial sequences within the nodule itself are difficult to assess. Nonetheless, several of the genera in the nodule microbiome analysis directly correspond to the nodule isolate genera.

**Figure 3.**
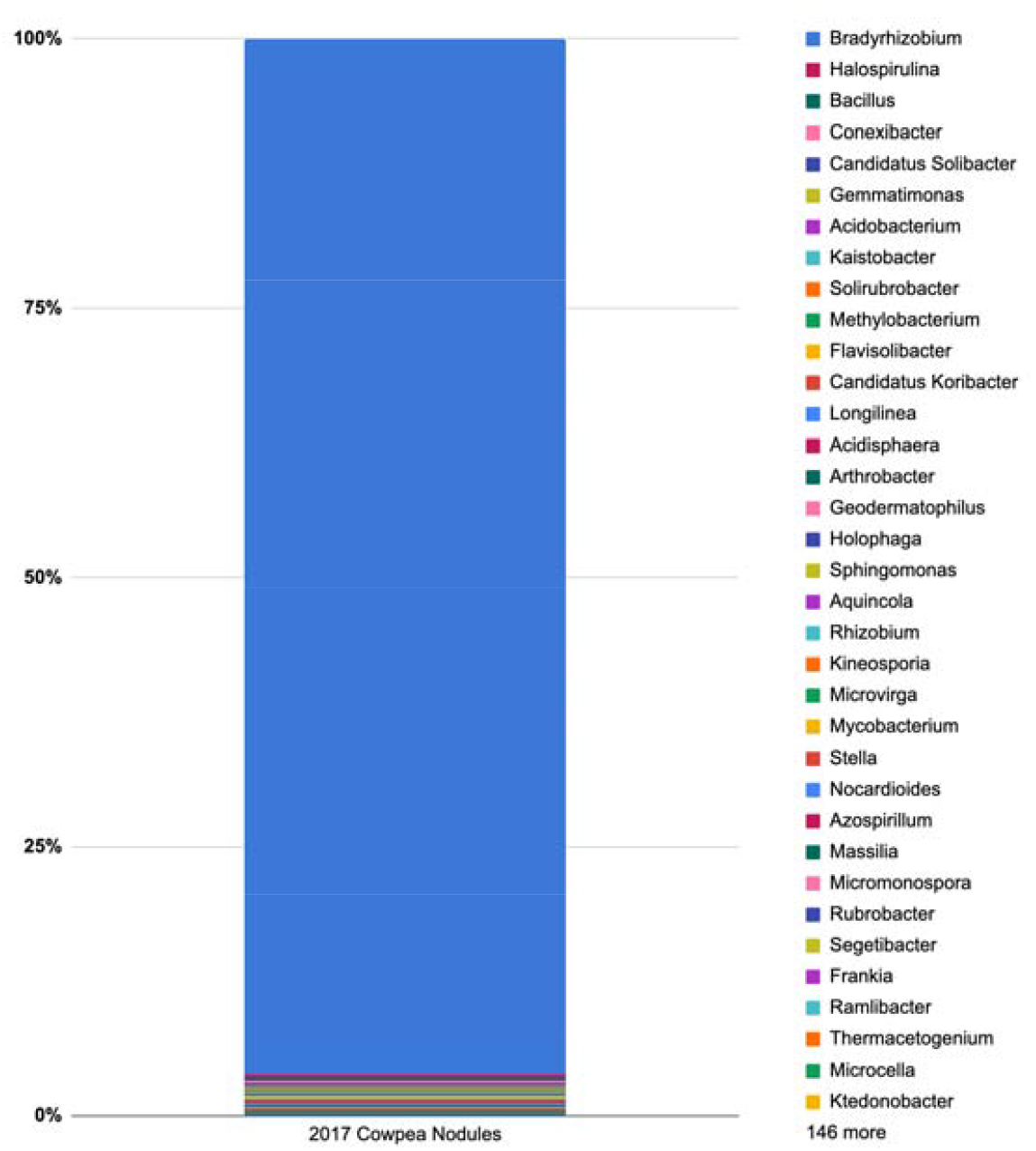
Cowpea nodule microbial community composition (by genus) as determined by analysis of environmental DNA. Cowpea plants were grown in the 2017 Botswana soil sample, and environmental DNA extraction and analysis was conducted in early 2019.

The bacterial population of nodules based on sequencing the isolated eDNA differs in terms of representation from the results obtained from isolating microbes from soil [28, 29]. The soil population is dominated by Actinobacteria, Proteobacteria, and Firmicutes (Fig. 1). Although a large number of Gram-positive species are detected in the soil microbiome analysis, they are detected at very low levels in the nodule microbiome (Fig. 3). Nevertheless, some genera such as *Bacillus* as well as actinomycetes, especially *Micromonospora*, are repeatedly isolated from nitrogen-fixing nodules and also detected in the nodule microbiome analysis. Coupled with the fact that several of these species, when inoculated with rhizobia frequently enhance the symbiosis [25, 30], this strongly suggests that they have a positive effect on the symbiosis.

The difference between the soil and nodule microbial populations in terms of numbers of microbes is reminiscent of the differences in the numbers of bacteria found in the rhizosphere versus the endosphere and between the rhizosphere and rhizoplane (surface) communities in other plant systems such as pepper [8] and maize [31]. Whether or not this is a specific selection for a large number of beneficial bacteria to protect the root or leaf surface from pathogen attack as suggested by the camouflage hypothesis [32] or that normally surface bacteria are excluded from internal tissues and only some of the bacteria that enter roots and nodules are “cheaters” [33] is difficult to determine at this time. In contrast, rhizobia are actively selected by the host plant for their symbiotic traits in response to active recognition between the host and its symbiont. Whether a similar recognition system operates between PGPB and plant surfaces is not known [34]. Although the mechanisms used by the non-rhizobial endophytes to enter the root and the nodule frequently involve the secretion of hydrolytic enzymes such as cellulase and pectinase, are these enzymes induced because the bacteria are recognized, and if so, what are the signals to which the endophytes are responding? To our knowledge, the mechanisms underlying how a coinoculation between rhizobia and PGPB triggers plant growth stimulation are not well understood.

## Conclusions

Traditional agriculture methods, which include legume intercropping, crop rotations with legumes, and use of legume cover crops, not only guaranteed that sufficient nitrogen was available for crops to be planted in the next growing season, but also ensured that the appropriate microbiome bacteria, particularly for legume cultivation, would be replenished for subsequent planting seasons. However, the practice of applying biofertilizers to seeds consisting of a single bacterial inoculant, which may or may not be competitive in modern agricultural soils, coupled with an inadequate knowledge of the microbial composition of different soils with or without crops as well as an over-dependence on N fertilizers, has exacerbated a loss of soil fertility. What is needed now is the development of optimal microbiomes that are matched to a particular crop and soil environment similar to the strategies of trying to enhance the health of humans by supplementing their either ill-nourished or antibiotic-affected microbiomes [35, 36]. Paying better attention to the evolution of the plant-bacteria symbiosis as well as their ecosystems [37], a highly specialized hologenome [38] will give us greater insight into the complex interaction between plants and their beneficial microbes, both of which live and interact together in a soil-microbial community.

To this end, we have isolated a large number of bacteria from soil collected from fallow farmland in Botswana, and also from root nodules of cowpea plants grown as “trap plants” in the same soil. We will continue to test these and other PGPB with or without rhizobial coinoculation for their effects on plant growth and responses. In addition, inoculants need to be rigorously screened to verify that they have no adverse effects on livestock or humans or on agricultural and indigenous plants. These are not trivial undertakings, and rigorous testing not only in the lab but also in the field must be performed. Currently, very few Botswana farmers use inoculants for their crops and indeed the inoculant industry is non-existent in Botswana and many parts of Africa. Inoculation with rhizobia itself is an uncommon practice. Instead, farmers rely on the indigenous soil microbes with the hopes that they will have a positive effect on crop growth following planting or they add N, P, and K fertilizers to their fields. Indeed, fertilizer consumption in Botswana has increased from 58.3 kilograms per hectare in 2014 from 89.6 kilograms per hectare in 2016, a 48.43% increase although fertilizer usage fluctuates from one year to the next [39]. Fertilizer addition definitely promotes plant growth, but its overuse also results in soil degradation due to loss of the indigenous soil microflora and fauna as well as pollution of waterways due to runoff. Farmers in Botswana and elsewhere need support for using inoculation technology and should be given incentives to do so. Achieving environmental balance is a daunting endeavor but using indigenous microbes will help us achieve the goal to grow crops sustainably while simultaneously restoring soil health.

## Supplementary Materials

Figure S1: Image of *Tephrosia purpurea*, Table S1. Characterization of bacteria isolated from 2017 and 2019 *Tephrosia purpurea* rhizosphere soil samples.

## Author Contributions

Conceptualization, AMH, FPM; methodology, PMH, EAH, KR, MK, JGI scientists; software, JGI scientists; validation, all authors; formal analysis, all authors; investigation, all authors.; resources, AMH, FPM, KR; data curation, AMH, KR; writing—original draft preparation, AMH, FPM, MK, EAH, PMH; writing—review and editing, AMH, FPM, MK, PMH, EAH; visualization, AMH, FPM; supervision, AMH, MM, JC, PY, EAH; project administration, AMH; funding acquisition, AMH. All authors have read and agreed to the published version of the manuscript.

## Acknowledgments

We are grateful to Dr. Erin Sanders-O’Leary, who encouraged us to pursue this research project in an undergraduate laboratory class setting. We acknowledge numerous undergraduate students who either were part of this UCLA laboratory course or were summer visitors in the Hirsch lab. We are especially grateful to the following students who contributed: C.A. Aliaga, M. Arrabit, D. Bagdasarian, M. Beyder, C. Bhagawat, P. Bruder, N. Bueno, A. Caceres, E. Curiel, G. Dogra, A. Eade, A. Fong, A. Haworth, D. Lee, T. Luong, P. Kim, T. Lui, T. Luong, S. Martikyan, S. Orellana, P. Patel, L. Poonnopatum, N. Saini, A. Sarquiz, D. Shia, R. Smith, G. Sunga, S. Surendranathan, C. Tran, D. Ullrich.

Seeds of *Lebeckia ambigua* were generously provided by John Howieson (Murdoch University, Perth, Australia), *Vigna subterraneana* by Felix Dakora (Tshwane University of Technology, Pretoria, South Africa), and *Macroptilium atropurpureum* by Heinz Hoben (BioNext, Wichita, KS, USA).

