## Supplementary material for "Isolation of potential plant growth-promoting bacteria from nodules of legumes grown in arid Botswana soil": Figure S1

**Supplementary FIGURE and Table.**

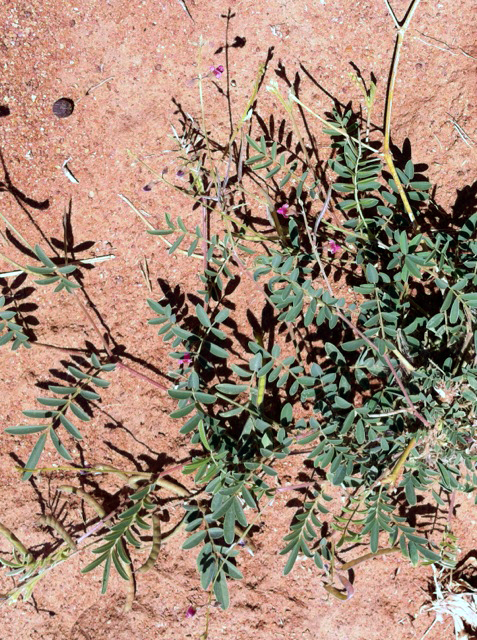

Fig. S1. *Tephrosia purpurea* growing in a farm field at Botswana University of Agricultural and Natural Resources.

**Table S1. List and characterization of bacteria isolated from 2017 and 2019 *Tephrosia purpurea* rhizosphere soil samples.**

| Sample | Closest Identity Match | CAS | PVK | Cellulase |
| --- | --- | --- | --- | --- |
| YMA-1 | *Arthrobacter* sp. | nd^1^ | + | - |
| R2A-2 | *Massilia yuzhufengensis* | nd | + | + |
| YMA-2 | *Microvirga zambiensis* | nd | + | + |
| LB-1 | *Paenibacillus cineris* | nd | + | + |
| LB-2 | *Bacillus drentensis* | nd | + | + |
| LB-3 | *Paenibacillus popilliae* | nd | + | - |
| LB-4 | *Bacillus safensis* | nd | + | + |
| L3 | *Bacillus wiedmannii* | + | - | - |
| L23 | *Bacillus amyloliquefaciens* | + | + | - |
| L31 | *Bacillus pumilus* | - | + | + |
| L44 | *Bacillus velezensis* | - | + | + |
| Y14 | *Bacillus aryabhattai* | - | + | + |
| Y27 | *Paenibacillus polymyxa* | + | + | + |
| Y30 | *Bacillus safensis* | - | + | + |
| Y34 | *Bacillus pumilus* | - | + | - |
| Y35 | *Brevibacillus brevis* | - | + | + |
| Y36 | *Kitasatospora cheerisanensis* | - | + | + |
| R36 | *Bacillus australimaris* | - | + | + |
| R61 | *Paenibacillus chitinolyticus* | - | + | - |
| L50 | *Bacillus subtilis* subsp. *inaquosorum* | + | + | + |
| R70 | *Streptomyces asterosporus* | + | + | + |
| NS TY 4 | *Domibacillus robiginosus* | nd | nd | nd |
| NS AG 1 | *Curtobacterium* sp. | nd | nd | nd |
| NS AG 2 | *Arthrobacter* sp. | nd | nd | nd |
| NS AG 4 | *Bacillus funiculus* | nd | nd | nd |
| NS AG 5 | *Bacillus niacini* | nd | nd | nd |
| NS AG 7 | *Staphylococcus sciuri* | nd | nd | nd |
| 2015 - 20 | *Oerskovia enterophila* YIM103440 | nd | nd | nd |
| 2015 - 32 | *Staphylococcus warneri* GS1 | nd | nd | nd |
| 2015 - 18 | *Bacillus altitudinis* GC315 | nd | nd | nd |
| 2015 - 5 | *Bacillus aryabhattai* GB61 | nd | nd | nd |
| 2015 - 3 | *Bacillus siamensis* MS41 | nd | nd | nd |
| 2015 - 9 | *Paenibacillus ehimensis* NPUST1 | nd | nd | nd |
| 2015 - 8 | *Paenibacillus elgii* JCK5075 | nd | nd | nd |
| 2015 - 15 | *Paenibacillus silvae* RR59 | nd | nd | nd |

^1^nd, not determined
